# Metagenomic discovery and co-infection of diverse wobbly possum disease viruses and novel hepaciviruses in Australian brushtail possums

**DOI:** 10.1101/795336

**Authors:** Wei-Shan Chang, John-Sebastian Eden, William J. Hartley, Mang Shi, Karrie Rose, Edward C. Holmes

**Affiliations:** Marie Bashir Institute for Infectious Diseases and Biosecurity, Charles Perkins Centre, School of Life and Environmental Sciences and Sydney Medical School, University of Sydney, Sydney, NSW, Australia; Westmead Institute for Medical Research, Centre for Virus Research, Westmead, NSW, Australia; Australian Registry of Wildlife Health, Taronga Conservation Society Australia, Mosman, NSW, Australia; James Cook University, College of Public Health, Medical & Veterinary Sciences, Townsville, QLD, Australia

**Keywords:** RNA-sequencing, meta-transcriptomics, wobbly possum disease, neurological disease, arterivirus

## Abstract

**Background:** Australian brushtail possums (*Trichosurus vulpecula*) are an introduced pest species in New Zealand, but native to Australia where they are protected for biodiversity conservation. Wobbly possum disease (WPD) is a fatal neurological disease of Australian brushtail possums described in New Zealand populations that has been associated with infection by the arterivirus (*Arteriviridae*) wobbly possum disease virus (WPDV-NZ). Clinically, WPD-infected possums present with chronic meningoencephalitis, choroiditis and multifocal neurological symptoms including ataxia, incoordination, and abnormal gait.

**Methods:** We conducted a retrospective investigation to characterise WPD in native Australian brushtail possums, and used a bulk meta-transcriptomic approach (i.e. total RNA-sequencing) to investigate its potential viral aetiology. PCR assays were developed for case diagnosis and full genome recovery in the face of extensive genetic variation.

**Results:** We identified a distinct lineage of arteriviruses from archival tissues of WPD-infected possums in Australia, termed wobbly possum disease virus AU1 and AU2. Phylogenetically, WPDV-AU1 and WPDV-AU2 shared only ∼70% nucleotide similarity to each other and the WPDV-NZ strain, suggestive of a relatively ancient divergence. Notably, we identified a novel and divergent hepacivirus (*Flaviviridae*) - the first in a marsupial - in both WPD-infected and uninfected possums, indicative of virus co-infection.

**Conclusions:** We have identified a distinctive marsupial-specific lineage of arteriviruses in mainland Australia that is genetically distinct from that in New Zealand, in some cases co-infecting animals with a novel hepacivirus. Our study provides new insight into the hidden genetic diversity of arteriviruses, the capacity for virus co-infection, and highlights the utility of meta-transcriptomics for disease investigation and surveillance in a One Health context.

## Background

Wildlife experience a diverse array of infectious diseases, many of which can impact animal welfare, biodiversity, environmental, livestock, and human health, tourism, and trade. While there is a growing volume of research into the ecology of wildlife infectious diseases, there is a marked absence of research on neglected host taxa, neglected geographic areas, and neglected wildlife pathogens, particularly those that do not threaten human or livestock health [1]. Similarly, there has been insufficient investigation and surveillance of “biodiversity diseases” where there are no perceived direct agriculture or human health links, including such high profile examples as chytrid fungus infection in amphibians and white nose syndrome in bats [2]. In our fast moving and highly connected world, rapid diagnosis and response are required to identify, understand and respond to emerging disease threats. Bias over the nature of emerging disease in wildlife, particularly before a diagnosis has been established, can rob us of the opportunity to conduct science-based risk assessments into potential collective health threats and impacts. Wildlife diseases, in particular, may have implications for the loss of natural biomass and important ecosystem services such as clean water and pollination. Wildlife diseases are also central to a One Health perspective because they can threaten native species with extinction [3], can be used as biological controls against invasive pest species, and may act as conduits for viruses to move to domestic species and ultimately humans [4].

Although common and widely distributed across Australia, the brushtail possum (*Trichosurus vulpecula*) is protected under conservation legislation. In contrast, the species is a highly successful invasive vertebrate pest in New Zealand, where it was introduced from Tasmania in the 1830s to build a local fur industry [5]. The species then rapidly adapted to the natural environment of New Zealand, causing devastating destruction to native forests and wildlife, and functioning as an important reservoir of the bacteria *Mycobacterium bovis* [6] and *Leptospira* sp. [7, 8]. Brushtail possums in New Zealand are estimated ted to cause $NZ35M/year in agricultural losses and the shared costs of attempted control exceed $100M/year [9].

Wobbly possum disease (WPD) is a severe neurological disease of Australian brushtail possums in New Zealand, caused by a virus of the family *Arteriviridae* (order *Nidovirales*) termed wobbly possum disease virus (WPDV) [10]. WPDV was first recognized in New Zealand in diseased possums present at a research facility in 1995, and has subsequently been found in a free-living population in the same country [11, 12]. In New Zealand, WPD presents as severe, multifocal neurological signs, including incoordination, loss of balance, head tilt, circling, difficulty climbing, abnormal gait, aimless wandering and can include central blindness. Most animals are found thin or emaciated and are either anorexic or exhibit abnormal behaviours such as daytime feeding [12-14]. There are no known treatments for WPD and affected animals presenting to wildlife carers most often die or are euthanized. Histologically, cases of WPD in New Zealand are similar to those from experimentally induced infection with WPDV and include non-suppurative meningoencephalitis, perivascular inflammation in multiple organs, most consistently liver and kidney, and salivary gland [15]. The disease in New Zealand has been investigated through multiple experimental infection trials, transmission trials and the development of multi-modal diagnostic tools including RT-qPCR, *in situ* hybridisation, immunohistochemistry, growth of the virus in possum macrophage cell culture, and indirect ELISA serology [10-12, 15, 16]. In contrast, WPD in brushtail possums in Australia has been sparsely described and investigated [17, 18].

Serological surveys in New Zealand from archival cases suggested an estimated WPD prevalence of 30% in the free-living possum population [19]. Sporadic cases and rare outbreaks of WPD have been reported in mainland Australia and Tasmania based on similar clinical presentations, histopathologic changes and exclusion of differential diagnoses such as trauma, bacterial infection, haemorrhage from rodenticide ingestion, *Angiostrongylus cantonensis* or *Toxoplasma gondii* infection. Previous investigations diagnosed WPD in 21 of 31(68%) brushtail possums with neurological dysfunction in the Sydney basin, New South Wales between October 1998 and June 2010 [18]. Although a viral aetiology for mainland Australian WPD has been suspected, a causative agent has not been established and there is considerable uncertainty surrounding the microbial causes, potential transmission routes, the infectious dose required to incite disease, carrier states, and other elements of host-virus ecology.

Arteriviruses tend to be host species-specific and cause a number of distinct disease syndromes with unique pathogenesis and epidemiology. Notable pathogens in this family include equine arteritis virus, porcine reproductive and respiratory syndrome virus, and simian haemorrhagic fever virus [20]. These viruses are phylogenetically distinct from the recently emerging *Nidovirales* from the family *Coronaviridae* that have been identified as pathogens of Australian reptiles, causing respiratory disease in shingleback lizards [21] and a mass mortality in a species of freshwater turtle associated with Bellinger River virus [22]. Phylogenetically, WPDV falls into what appears to be a divergent branch lineage of the *Arteriviridae* [23]. Current hypotheses for the origin of WPDV in New Zealand are that the virus evolved and emerged in possums through cross-species transmission after translocation, or the virus is possum-specific and was carried by animals during their introduction from Australia [5]. The identification of WPDV in native animals on the Australian mainland would be important evidence supporting the latter.

Hepaciviruses are a genus of positive-sense, single-stranded RNA viruses from the family *Flaviviridae*, the best characterised of which is Hepatitis C virus (HCV) due to its association with hepatitis and hepatocellular carcinoma in humans worldwide [24]. However, other hepaciviruses have been identified in dogs [25], horses [26], rodents [27], bats [28] and ducks [29], and new hepaciviruses are regularly being described in a variety of animal species. To date, however, hepaciviruses have not been reported from any marsupial species.

Herein, we review archived Australian cases of WPD to build a more robust syndrome description and, where frozen tissues were available, applied total RNA sequencing (“meta-transcriptomics”) to determine the infectious aetiology and potential origins of WPD in mainland Australian possums, particularly in comparison with syndromes and agents previously identified in animals from New Zealand. Accordingly, this study provides new insights into the diversity and origins of a novel group of viral pathogens and illustrates the utility of meta-transcriptomics as a means to identify complex infections in a One Health perspective.

## Methods

### Sample collection

Samples were collected between May 1999 and June 2010 from nine brushtail possums found within the greater Sydney basin. All animals were handled under a series of NSW Office of Environment and Heritage Licences to Rehabilitate Injured, Sick or Orphaned Protected Wildlife (#MWL000100542). Samples were collected from affected possums post mortem, immediately after euthanasia via intravenous barbiturates delivered while the animals were under gaseous anaesthetic (2% isofluorane in 1L/min Oxygen - Isofluorane 100%, Zoetis, Australia) for veterinary examination. Fresh brain, liver and kidney were collected aseptically and frozen at −80°C. A range of tissues was fixed in 10% neutral buffered formalin, processed in ethanol, embedded with paraffin blocks, sectioned, stained with haematoxylin and eosin and permanently mounted with a cover slip. Samples were collected under the Opportunistic Sample Collection Program of the Taronga Animal Ethics Committee, and under scientific licences #SL10469 and SL100104 issued by the NSW Office of Environment and Heritage.

### Historical Case Review

All brushtail possum cases within the Australian Registry of Wildlife Health since systematic record keeping began in 1981 were evaluated for signalment, clinical signs and histological lesions. Animals with a combination of visual impairment, severe depression or central nervous system disturbance in conjunction with non-suppurative lesions in the central nervous system, optic tracts or eyes were considered to fit the WPD syndrome description. The severity of non-suppurative lesions in the meninges, cerebral cortex, brainstem, cerebellum, liver, kidney and eyes were graded on a scale of 0-4 where 0 represented no discernible lesions and 4 represented severe and extensive lymphoplasmacytic inflammation. Necrosis was graded on a scale of 0-4 where 1 represented mild, multifocal single cell necrosis and 4 characterised extensive malacia. Retinal atrophy was similarly assessed on a scale of 0-4 with grade 4 signifying a nearly aceullular ganglion cell layer.

### Pathogen discovery using meta-transcriptomics

For RNA extraction, archival tissues including brain, liver and kidney samples of animals were processed using the RNeasy Plus Mini Kit (Qiagen, Germany). RNA concentration and integrity were measured using a NanoDrop spectrophotometer (ThermoFisher Scientific, USA) and TapeStation (Agilent). Samples were then pooled in equal proportions based on tissue type and/or individual cases for different purposes (Table 1). Illumina TruSeq stranded RNA libraries were prepared on the pooled samples following rRNA depletion using the RiboZero Gold kit (Epidemiology). Paired-end (100 bp) sequencing of the rRNA-depleted RNA libraries were then performed on an Illumina HiSeq 2500 system at the Australian Genome Research Facility (AGRF), Melbourne.

The meta-transcriptomic analytical pipeline was constructed based on the methods previously used in our group [30-32]. Accordingly, RNA sequencing reads were trimmed of low quality bases and adapter sequences and *de novo* assembled using Trinity 2.1.1 [33]. Assembled sequence contigs were annotated using both nucleotide and protein BLAST searches against the NCBI non-redundant sequence database. To identify low abundance organisms, the sequence reads were also annotated directly using a Diamond Blastx search against the NCBI virus RefSeq viral protein database (with an e-value cutoff of <10^−5^). Open reading frames were then predicted from the viral contigs in Geneious v11.1.2 [34] with gene annotation and functional predictions made against the Conserved domain databases (CDD) [35] and Geneious v11.1.2. Virus read abundance was assessed using a read mapping approach available in the BBmap program [36].

### PCR assays and genome sequencing of WPDV and possum hepacivirus

Our metagenomic analysis suggested the presence of both WPDV and a novel hepacivirus (see Results). To confirm their presence in each archival tissue from both diseased or healthy possums, the initial PCR primers were designed based on aligned sequence reads generated from RNA-seq libraries (Table 1). Subsequently, SuperScript IV VILO cDNA synthesis system (Invitrogen) was used to reverse transcribe the RNA from individual cases. The cDNA generated from the sampled tissues was used for viral specific PCRs targeting regions identified by RNA-Seq. All PCR assays were performed using Platinum SuperFi DNA polymerase (Invitrogen) with a final concentration of 0.2 μM for both forward and reverse primers.

Long, overlapping PCR assays were also developed to complete the genome of two representative Australian WPDVs (Table S1). Similarly, additional sets of PCR primers for hepacivirus genome sequencing were designed from RNA-sequencing data of individual affected cases (Table 1). All PCR products were visualized using agarose gel electrophoresis and confirmed by both Sanger and MiSeq sequencing.

### Wobbly possum disease virus qRT-PCR

A quantitative RT-PCR assay targeting the conserved RdRp domain was developed to identify viruses of all three different lineages of WPDVs (New Zealand, Australia 1 and 2). The qRT-PCR primers and probe sequences (Integrated DNA Technologies, USA) were designed as described in SI Table 1. PrimeTime Gene Expression Mastermix (Integrated DNA Technologies, USA) was used to perform the assays. For absolute quantification, the pooled genome amplicons were prepared as standards across aerial 10-fold dilution series of known quantities and used to determine virus copy number.

### Phylogenetic analysis

To determine the evolutionary relationships of the WDPVs and the novel hepacivirus identified in this study, we performed phylogenetic analyses of amino acid sequences of the NSP2 protein (the RNA-dependent RNA polymerase (RdRp) of arteriviruses) and the complete polyprotein of hepacivirus, respectively. All sequences were aligned using MAFFT version 7 [37] with the L-INS-i algorithm, with all ambiguously aligned regions removed using TrimAL (v.1.4.1) [38]. Maximum likelihood trees of both data sets were then estimated using IQ-TREE 1.6.7 [39] utilising the best-fit model of amino acid substitution (LG+F+I+ Γ 4). Statistical support for individual nodes was assessed using a bootstrap approach with 1000 replicates, and all trees were midpoint rooted. Finally, phylogenetic trees were visualized using FigTree v1.4.3 [40].

### Nucleotide Sequence Accession Numbers

The RNA sequencing data generated in this study have been deposited in the GenBank Sequence Read Archive under accession numbers PRJNAXXXX. All consensus genome sequences of identified viruses have been uploaded in GenBank under accession numbers XXXXX to XXXXXX.

## Results

### Clinical and histological description of Australian wobbly possum disease virus cases

Signalment, gross and histopathology data were collated from 474 brushtail possums, 49 of which fit the syndrome description for WPD. A table chronologically summarising the clinical and histological findings of WPD in brushtail possums in Australia is included as Table S2. The index case of WPD on mainland Australia was a juvenile male possum found in Mosman, NSW in December 1983, while the index case in Tasmania was November 1985 based on Registry records. The majority of WPD cases occurred in adult female possums (35 female, 9 male, 6 unknown sex, 42 adult, 3 juvenile, 4 unknown age).

Blindness was the most common presenting sign (n=35) and was characterised by dilated and unresponsive pupils and lack of a menace reflex. Depression or docility were noted in six animals. A further 4 animals were found moribund. Clinical signs attributable to the nervous system included ataxia (n=9), circling (n=4), strabismus (n=1), paralysis (n=2), nystagmus (n=1), and knuckling (n=1), and these animals were most often concurrently blind. Some WPD affected animals were maintained in care for many weeks, with stable or slowly progressive signs. Ophthalmological examination of blind possums often revealed an optic disc that was pale and lacked the normal fundus vascular tuft. The body condition of affected possums ranged from excellent to emaciated. The most common cause of death was euthanasia.

Common microscopic lesions in WPD affected possums are illustrated in Figure 1 and included mild to severe non-suppurative leptomeningitis, multifocally extending into the perivascular spaces of the brainstem, cerebrum and cerebellum. Cellular infiltrates ranged from 1 to 12 cells deep, and were composed of lymphocytes, plasma cells and smaller numbers of macrophages. Small foci of gliosis were multifocally evident in the neuropil of severely affected animals. Neutrophils were seen among the inflammatory infiltrates in only two animals. Necrosis was a less consistent finding and ranged between mild, multifocal single cell necrosis of nerve cell bodies, particularly affecting cerebellar Purkinje cells, to foci of malacia or cerebellar folial atrophy. Wallerian degeneration and spongiotic change were observed in the optic tracts of the brain, and optic nerve, often in association with non-suppurative inflammation in the optic nerve, perineurium, retrobulbar fat, and sclera. Ocular lesions included mononuclear cell infiltrates, as above, evident variably within the choroid, retina, ciliary body and iris. Retinal atrophy was evident as loss of cellularity within the ganglion cell and nuclear layers, and occasionally a loss of rods and cones. Lymphoplasmacytic infiltrates in renal and hepatic tissue were only evident within animals originating from Tasmania, except in rare cases where concurrent disease could account for lesions.

**Figure 1.**
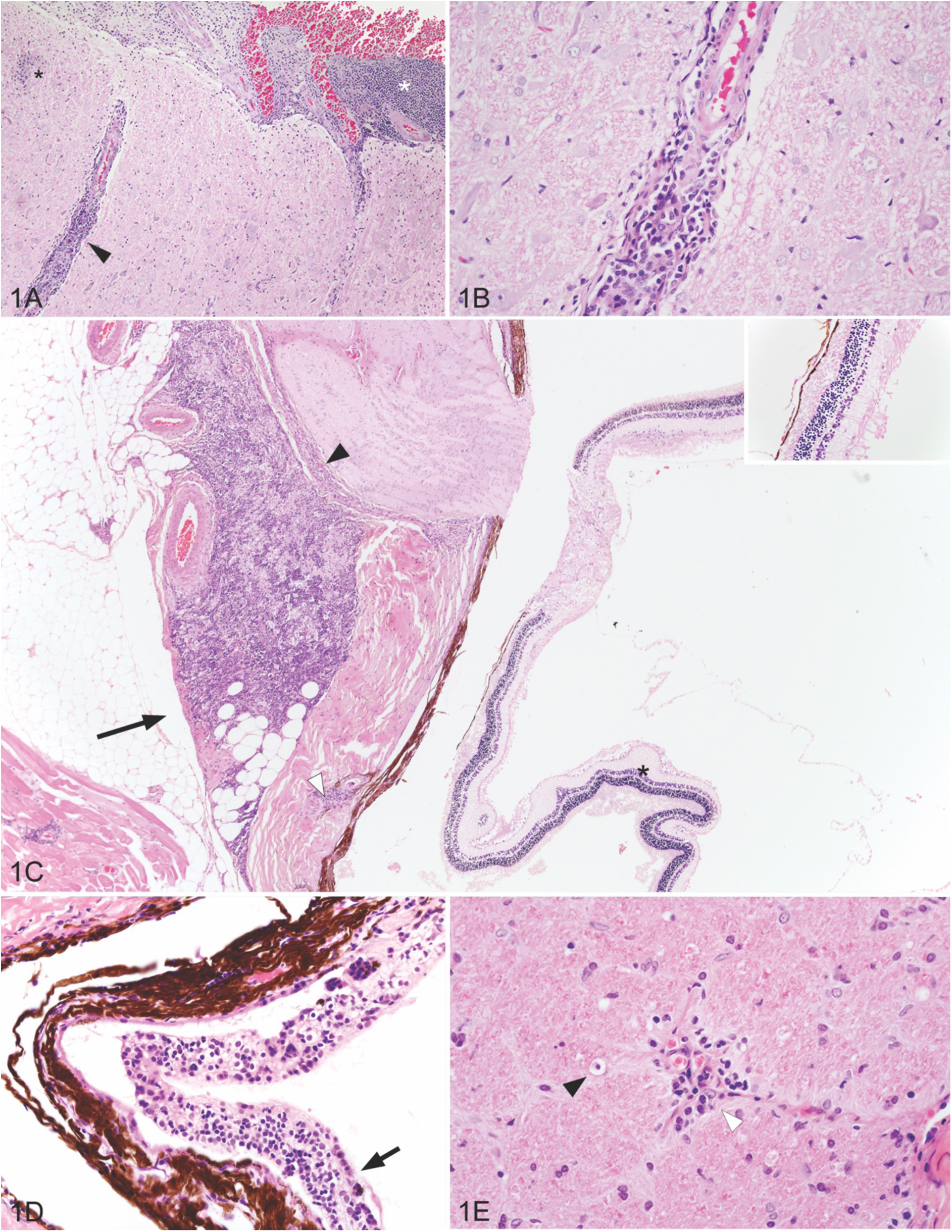
Histopathologic findings in Australian wobbly possum disease cases (hematoxylin and eosin). (A) Focal gliosis (black asterix), non-suppurative cellular infiltrates distending the leptomeninges (white asterix), and surrounding blood vessels (black arrowhead) within the brainstem. (B) Lymphocytes, plasma cells and scattered macrophages around a neural blood vessel. (C) Non-suppurative infiltrates in retrobulbar adipose tissue (black arrow), the perineurium of the optic nerve (black arrow head), and the scleral perivascular space (white arrow head). Retinal atrophy with an acellular ganglion cell layer (black asterix and inset). (D) Distention of the choroid layer with non-suppurative inflammation (arrow). (E) Optic nerve with Wallerian degeneration, illustrated by a macrophage within an axonal chamber (black arrowhead), and a non-suppurative perivascular infiltrate (white arrowhead).

### Meta-transcriptomic discovery of novel WPDV and hepacivirus in Australia brushtail possums

In total, eight rRNA-depleted RNA sequencing libraries were constructed, generating reads ranging from 23,480,309 to 26,509,764 per pool (Table 1). The first (Pool 1) contained the pooled brain RNA from three suspected WPDV-infected possums. Our initial analysis did not identify any assembled viral contigs including any related to WPDV or other arteriviruses. As low abundance pathogens may not have sufficient coverage for assembly, the trimmed sequence reads were then further annotated directly (i.e. without assembly) against the NCBI NR database using Diamond BlastX. This analysis revealed six sequence reads (three sequence pairs) with homology to WPDV from New Zealand (GenBank accession: NC_026811). In addition to the WPDV reads, eight sequence reads from a novel hepacivirus were present in the same library. Further RNA sequencing libraries were then prepared from the three individual cases found to be positive for WPDV (of which all were co-infected with the hepacivirus) that included libraries from multiple tissues where available (Table 1: cases 3619, 7613, 2545). To determine if any other organisms were present in the remaining cases, we also sequenced a hepacivirus monoinfection (Table 1: case 2345) as well as a pool of RNAs from those cases negative for both WPDV and hepacivirus (Table 1: pool 2). Overall, the read abundance of WPDV and hepaciviruses in each library was low, ranging between 0.000343%-0.000007% (Table 1).

### Identification of divergent lineages of WPDV

Individual PCR assays were designed based on the each of the three arterivirus-like sequence pairs identified from the initial RNA library and used to confirm the presence of WPD-like virus in specific cases. However, these PCR assays selectively amplified the viruses from the WPD-affected cases suggesting extensive genetic variation and the presence of different lineages of WPDVs in Australia. Accordingly, a more sensitive qRT-PCR assay was developed based on the conserved RdRp domain and specifically designed to targeting all WPDVs in this study including the NZ prototype strain. WPDV sequences were confirmed by qRT-PCR for the three WPD-affected possums including individual brain, liver and kidney tissues. Importantly, this pan-WPDV qRT-PCR assay did not identify any new cases and the tissues tested from the control possums (including case 1-4) were all negative. From this, we were also able to determine viral loads, which for WPD-positive tissues ranged from 21 to 4,777 copies per pg RNA and varied according to different tissue types. The highest level of WPDV expression was observed in liver tissues from case 2525 (4,777 copies/pg RNA) and from case 3619 (4,125 copies/pg RNA), followed by kidney (570 - 1,830 copies/pg RNA) and brain from each diseased possum (17 - 71 copies/pg RNA).

### Comparative genomics and phylogenetics of WPDVs

PCR and sequencing of the WPDV replicase gene showed that the three individual possums carried distinctive (i.e. case-specific) nucleotide variation in WPDV, comprising two distinct lineages (Figure 2). In addition, the WPDVs from different organs of each individual animal ormed discrete clusters, strongly suggesting that these results are not due to cross-sample contamination. Strikingly, pairwise alignment of the partial replicase polyprotein 1ab nucleotide genome (291 bp) revealed that the two lineages of Australian WPDV strains exhibited almost the same level of sequence divergence between them as they did to the New Zealand strain (∼70% nucleotide sequence similarity and 77% amino acid sequence similarity), indicative of substantial intra-specific virus diversity.

**Figure 2.**
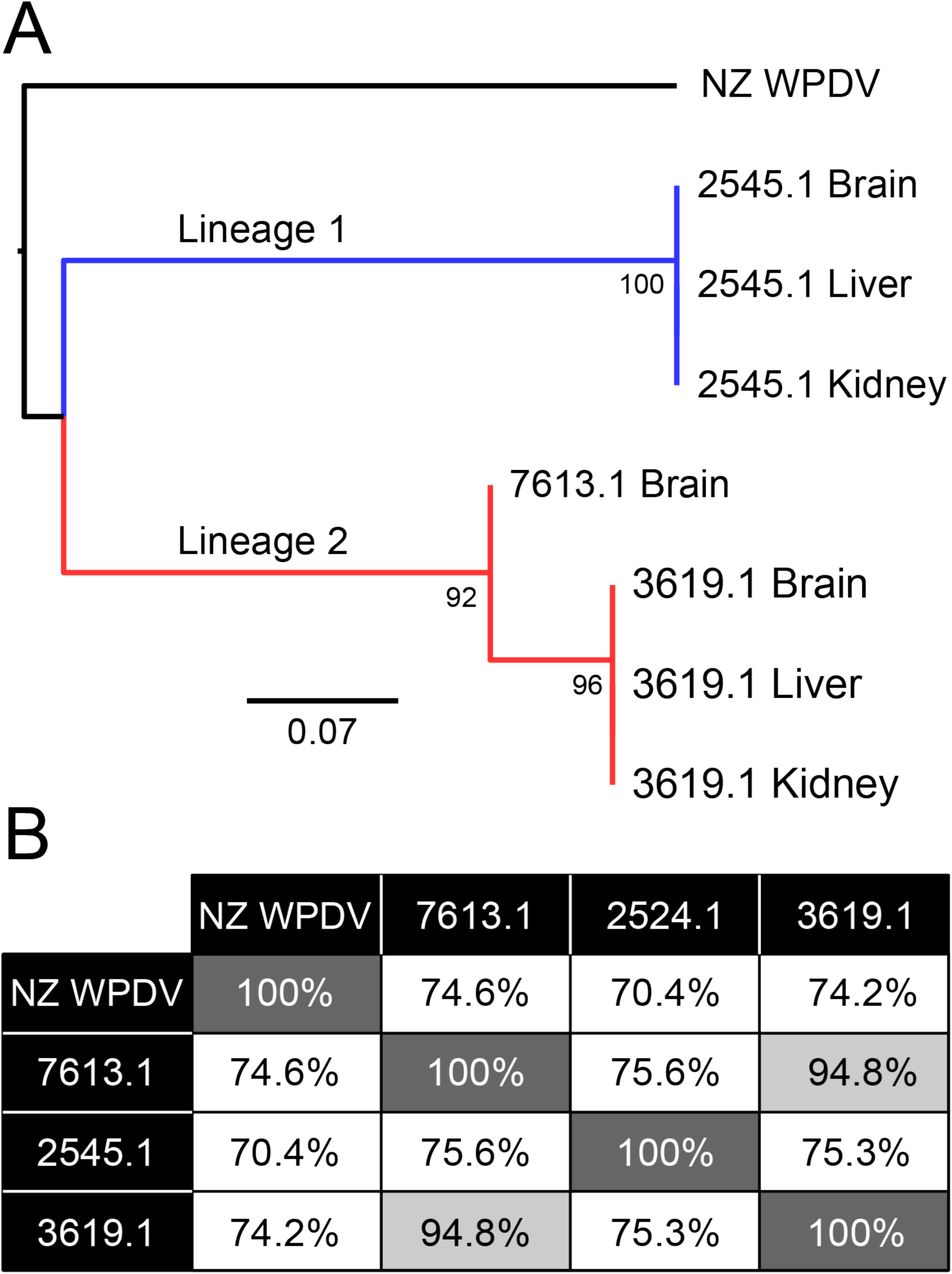
Identification of distinct lineages of WPDV in Australia. (A) Phylogenetic analysis of partial replicase polyprotein 1ab region of WPDV nucleotide genome (291 bp), showing the NZ reference strains compared to the three Australian cases identified here. Two major lineages of the Australian viruses were identified - denoted Lineages 1 and 2 - and coloured in blue and red, respectively. Nodes show bootstrap values from 1,000 replicates. The scale bar shows the number of nucleotide substitutions per site. (B) Pairwise comparisons of nucleotide similarity (%) across the viral genome.

Long overlapping PCRs were used to determine the near complete genomes of two representative WPDV strains, here termed WPDV-AU1 (case 2545) and AU2 (case 3619). In comparison to the WPDV-NZ strain, the two WPDV-AU genomes retained the arterivirus-like features, encoding a replicase polyprotein ORF1ab, glycoprotein (GP) 2, 3, 4, 5, and membrane protein (MP) and nucleocapsid protein (NP) (Figure 3A). The data generated here enabled us to determine the phylogenetic position of WPDV (both the Australian and New Zealand strain) within the *Arteriviridae* as a whole through a phylogenetic analysis of 1787 amino acids of the RdRp protein. Notably, WPDV, derived from a marsupial, fell as a sister-group to those arteriviruses previously documented in placental mammals (Figure 3B). In addition, the overall phylogeny of the *Arteriviridae* generally mirrors that of the vertebrate hosts from which they were sampled, with those viruses sampled from fish falling in the basal position, indicative of virus-host co-divergence for the entire evolutionary history of vertebrates.

**Figure 3.**
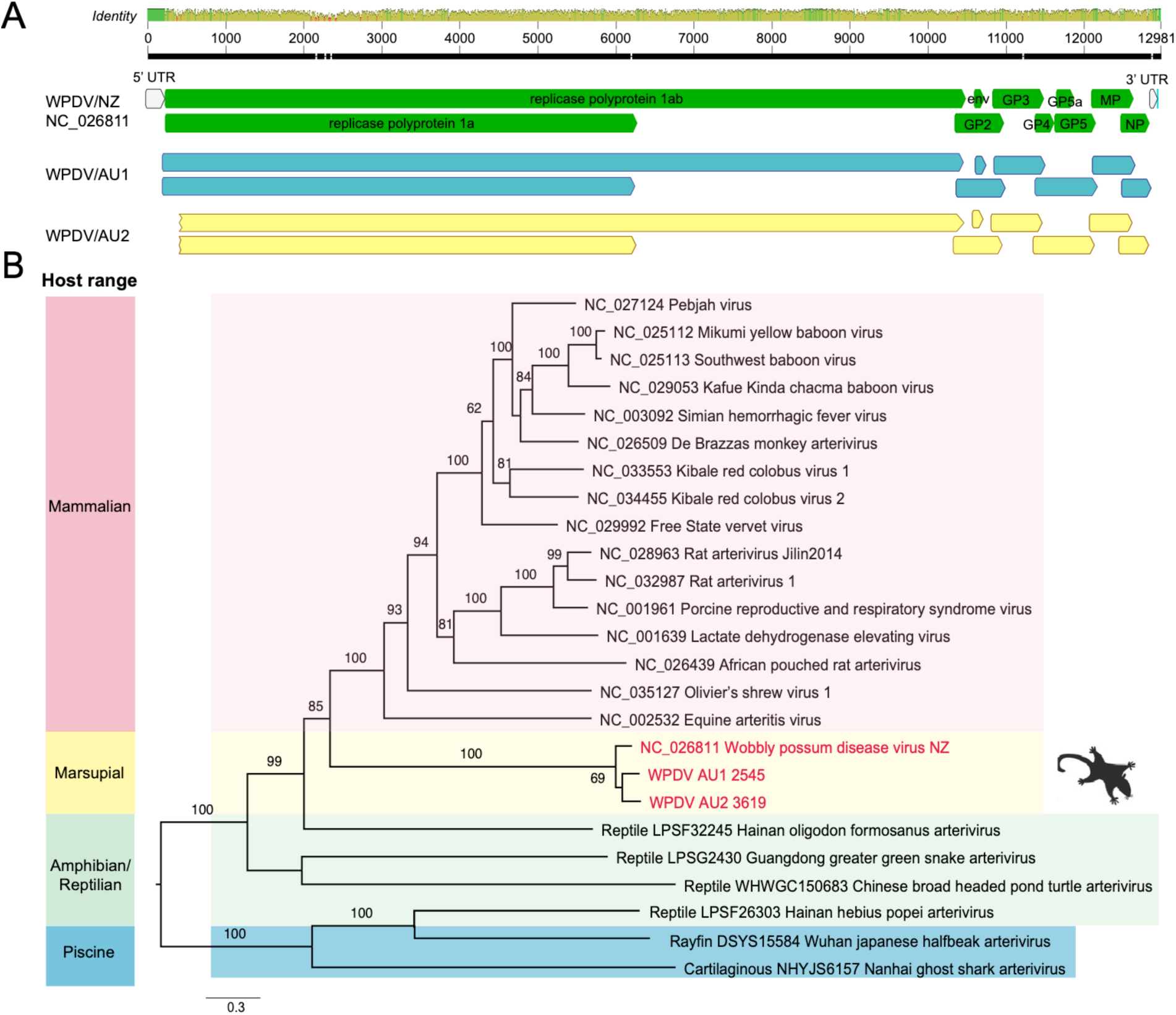
Genome organisation and evolutionary relationships of WPDV to other arteriviruses. (A) Genome features and comparison of three lineages of WPDV - WPDV-AU1, WPDV-AU2 and WPDV-NZ (GenBank accession: NC_026811). (B) Phylogenetic analysis of the arterivirus RdRp protein. Nodes show bootstrap values obtained from 1,000 replicates. Scale bar shows the number of amino acid substitutions per site. The tree was midpoint rooted for clarity only.

### A novel possum hepacivirus

Interestingly, our meta-transcriptomic analysis also identified a novel hepacivirus, that we have termed possum hepacivirus. This virus retains the classical hepacivirus features of a single-stranded, positive-sense RNA genome of 9,296 bp in length that encodes a single polyprotein. By further hepacivirus-specific PCR confirmation, we revealed the presence of the virus in liver, brain and kidney from three WPDV-affected possums and another two non-WPDV detected possums in our study, thereby revealing virus co-infection in animals with WPD (Table 2). A phylogenetic analysis of the complete polyprotein of hepacivirus (3260 amino acids) revealed that possum hepacivirus was most closely related to Norway rat hepacivirus 2 (accession number: YP_009109558.1) identified from brown rats (*Rattus norvegicus)* in New York City, US, with which it shared 41.2% sequence similarity in the polyprotein (Figure 4).

**Figure 4.**
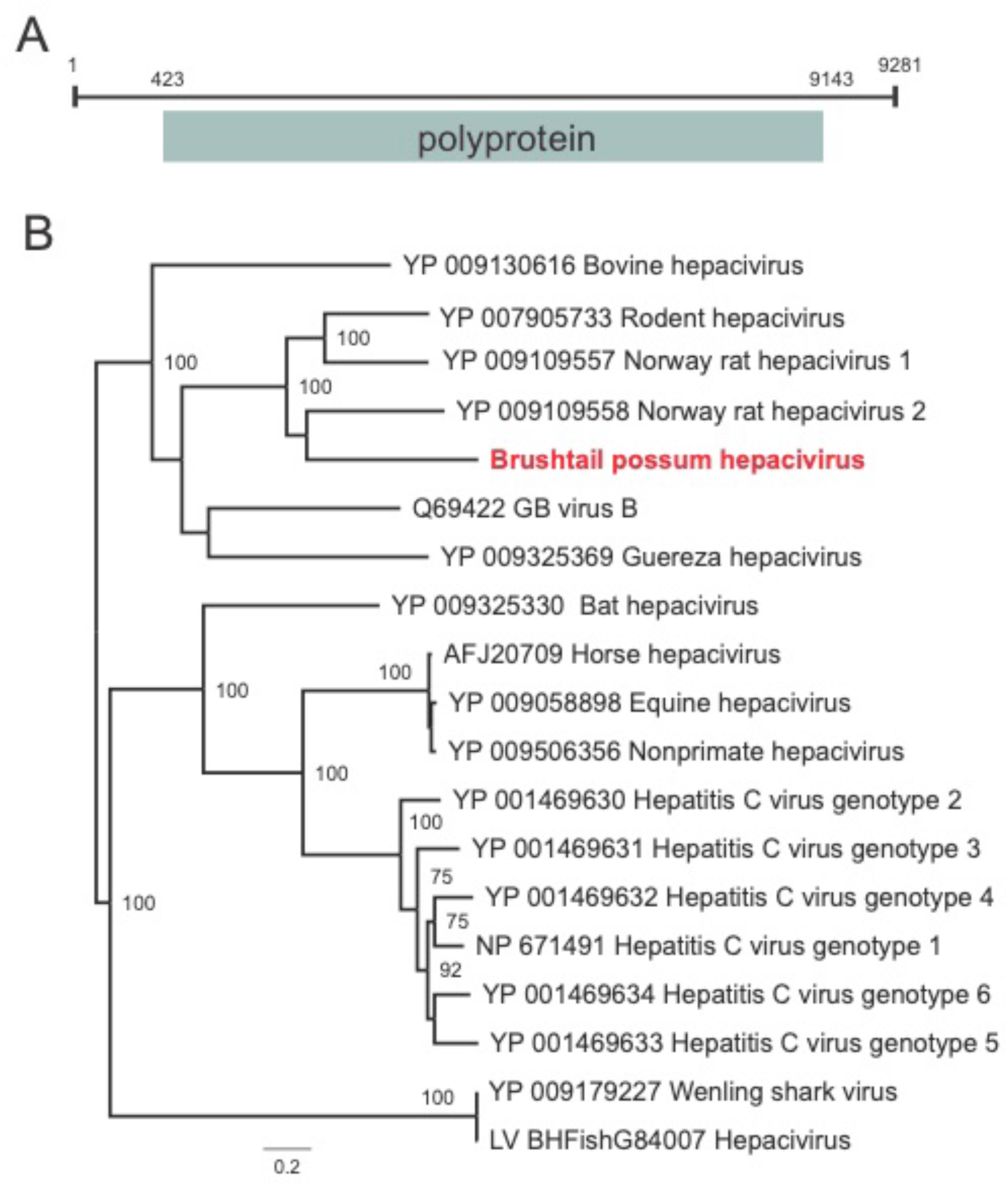
Evolutionary relationships of the novel hepacivirus identified from brush-tailed possums. Phylogenetic analysis of the polyprotein gene (RdRp protein) of hepaciviruses. The novel Brushtail possum hepacivirus is shown in red. Bootstrap values > 70% were presented for key nodes (1,000 replicates). The tree was midpoint rooted for clarity only. The scale bar shows the number of amino acid substitutions per site.

## Discussion

Our retrospective investigation and meta-transcriptomic analysis with subsequent PCR confirmation revealed that brush-tailed possums in mainland Australia that presented with a disease syndrome similar to that of WPD seen in New Zealand were infected with an arterivirus related to WPDV. In addition, three of the five animals presenting with WPD had evidence of co-infection with a novel hepacivirus, although whether this contributed to the symptoms and lesions observed is unclear.

Previously, strong evidence for the association between WPD and a viral aetiology was obtained using primary cell culture system for virus isolation [16], WPD-specific PCR assays and *in situ* hybridisation[11, 41]. By reproducing WPD in healthy possums with purified WPDV isolate inoculation, the histological changes highly met the natural infection status, fulfilling Koch’s postulates of causation. In addition, the establishment of the RT-qPCR and indirect ELISA assays for molecular and serological survey assists understanding disease prevalence in the wild population in New Zealand [41].

While the clinical and histological presentation of WPD as described here and by others varies geographically in wild possum populations in Australia, no active monitoring scheme is in place to explore the prevalence of the syndrome and diversity of associated viruses. Histologically, WPD in historical cases from Tasmania consistently had non-suppurative hepatitis and nephritis, similar to that described in the New Zealand form of WPD, and distinct from mainland Australian cases, which lack that feature. Importantly, we have designed a sensitive qRT-PCR assay that is able to detect divergent lineages of WPDV, including, WPDV-AU1, AU2, which should merit broader application in epidemiological studies of viral and syndrome prevalence. In addition, our results clearly show how a metagenomic approach is able to detect viral pathogens even at very low levels of read abundance, which will be applicable to other wildlife species and One Health scenarios.

Previous work [15] revealed that the viral loads of WPDV were significantly higher from the liver of experimental infected possums than other organs including kidney, brain, salivary glands and bladder, which is consistent with our qRT-PCR. Of note, these results also show consistently that the lowest viral loads were in the brain tissues. This was also apparent in the WPDV affected case 7614, which contained no detectable WPDV reads in the brain tissue by RNA-seq, but was clearly WPDV positive by PCR and Sanger assays. Although the qRT-PCR assays in our study successfully detected various strains, the viral loads among different tissues were generally lower than reference results from experimentally infected possums in the New Zealand studies [41]. This finding may reflect the often chronic nature of infection in diseased Australian possums, which were frequently examined several weeks after the onset of disease, but is also likely explained by the archival nature of our samples and hence relatively low preservation might affect the RNA integrity due to degradation. However, since our testing number is comparatively small, speculation on the overall distribution of WPDVs in Australian possum populations remains challenging. The collection of additional samples from different geographical locations, closely related, and sympatric species, larger serological and molecular surveys, in conjunction with population and ecological information will clearly assist understanding virus variation and transmission dynamics in the wild.

Our phylogenetic analysis reveals that WPDV formed a distinct lineage and that there has likely been long-term co-divergence between arteriviruses and their vertebrate hosts over many millions of years. As it is also clear that our sampling of the *Arteriviridae* is sparse, it is inevitable that more will be discovered through expanded metagenomic studies of diverse vertebrates. More challenging will be determining exactly how WPDV became established in brush-tailed possums in both Australia and New Zealand. Reports indicated that WPDV, or a virus causing similar neurological symptoms, has been present in free-living possum populations in New Zealand as far back as 1999 [12]. Our investigations track WPD cases back to 1983 and 1985 on mainland Australia and Tasmania respectively. As no evolutionary rate is available for WPDV it is currently impossible to perform a direct estimation of divergence times. However, the substantial sequence divergence between the Australian mainland and New Zealand strains suggests that their separation is relatively old and may have occurred close to when possums were introduced from Tasmania into New Zealand in the 1830s. Additional testing of Tasmanian brushtail possums will be an integral to addressing the question of whether WPDV was translocated to New Zealand with possums, or emerged and evolved afterwards.

A particularly notable aspect of this study was the presence of a novel possum hepacivirus also identified in our WPD-affected and non WPDV-detected possums. Most animal hepaciviruses are associated with chronic infection and strong hepatotropism, leading to hepatitis, cirrhosis, and severe hepatopathy [24]. More recently, a divergent Wenling shark virus (WLSV) hepacivirus was discovered in the liver of grateful catsharks (*Proscyllium habereri*) using the same bulk meta-transcriptomic approach as utilized here [42]. As the first documented hepacivirus identified from a marsupial, the novel Brushtail possum hepacivirus identified here expands the host range of this important group of viruses and again highlights the potential missing diversity of genus *Hepacivirus*. In addition, that the Brushtail possum hepacivirus falls in a clade of rodent hepaciviruses indicates that cross-species transmission has played a key role in shaping phylogenetic patterns. Clearly, natural infection routes and pathogenesis of this virus in possums merits additional work. Whether the novel possum hepacivirus identified here contributes to clinical disease or reduced fitness, alone or in conjunction with WPDV, and the status of both agents in wild populations remains largely unexplored, but is clearly a key area for future study including the presence in New Zealand animals or any of the inoculums used from previous challenge studies. More generally, an enhanced understanding of the roles of these and closely related organisms will shed important light on virome of non-eutherian mammals.

## Conclusions

The total RNA sequencing, or meta-transcriptomic, approach described has elucidated the pathogen likely responsible for a disease syndrome first detected in Australian mainland wildlife 36 years ago. Factors that may have contributed to this diagnostic delay and difficulty include the presentation in a common species, the sporadic rather than outbreak-based emergence of the syndrome, and the potential pre-conception that the disease posed no threat to human or livestock health. Nonetheless, the meta-transcriptomic technique described is becoming more rapid and cost-effective, and has demonstrated its capacity to remove many barriers delaying or obstructing traditional wildlife disease investigation. Additional benefits include its capacity to identify co-infections, and to detect and characterise non-cultivable microbes and those that diverge significantly from nearest phylogenetic neighbours. Data and knowledge generated from this technique will inform risk assessments addressing the potential threats and impacts of emergent pathogens in a One Health paradigm.

## List of Abbreviations

NCBI: National Center for Biotechnology Information.
NZ: New Zealand.
RT-PCR: Reverse transcription polymerase chain reaction.
WPD: Wobbly possum disease.
WPDV: Wobbly possum disease virus.

## Declarations

### Ethics approval and consent to participate

Samples were collected under the Opportunistic Sample Collection Program of the Taronga Animal Ethics Committee, and under scientific licences #SL10469 and SL100104 issued by the NSW Office of Environment and Heritage.

### Consent for publication

With the exception of WJH (deceased), all authors have agreed to submission of the final version of the manuscript.

### Competing interests

None declared.

### Funding

ECH is supported by an ARC Australian Laureate Fellowship (FL170100022).

### Authors’ contributions

Conceived the project - JSE, KR, ECH; collected samples - WJH, KR; performed laboratory work - WSC, JSE, KR; analysed the data - WSC, JSE, MS, KR; wrote the paper - WSC, JSE, MS, KR, ECH.

## Acknowledgements

This research was supported by Taronga Conservation Society Australia, Taronga Conservation Science Initiative, and New South Wales National Parks and Wildlife Service. Our thanks are extended to Jane Hall for diagnostic support and data management, Dr. Hannah Bender for graphic design of the photomicrographic plate, and to Drs. Rod Reece, Cheryl Sangster, Shannon Donahoe, Richard Montali, Cathy Shilton, Lydia Tong, Dr. Andrew Davis and Dr. David Obendorf for their diagnostic contributions on individual cases.

## Supplementary Information

**Table S1.** PCR primers and qPCR primers for WPDV used in this study.

**Table S2.** Presentation and pathology of Wobbly Possum Disease in brushtail possums in Australia. This table chronologically summarises the clinical signs and histological findings in brushtail possums considered to fit the syndrome description for Wobbly Possum Disease. Most possums originate from New South Wales on mainland Australia, except for those denoted with a ^T^ that originated from Tasmania.

